# A new long-acting GLP-1 derivative 6- KTP ameliorates body weight and lipid metabolism in DIO mice

**DOI:** 10.1101/846717

**Authors:** Peixiu Wang, Yanhong Ran

## Abstract

As a global epidemic, obesity has become the biggest challenge facing global and public health. Glucagon-like peptide-1(GLP-1) has been able to inhibit appetite, slow gastric emptying and reduce body weight. We designed a novel long-acting GLP-1 derivative 6-KTP based on wild-type GLP-1 and performed its pharmacodynamics study on obesity in DIO mice. DIO mice were treated once daily with subcutaneous injections of 6-KTP (1.8 mg/kg body weight), Liraglutide (0.4 mg/kg body weight) or vehicle (phosphate buffered saline (PBS), pH 7.4) for 12 weeks. The results show that 6-KTP decreased food intake, induced anorexia and weight loss, and improved blood lipid and lipid metabolism in DIO mice.

## Introduction

Obesity is associated with morbidity and mortality of chronic diseases and other health problems, including cardiovascular disease, type II diabetes, musculoskeletal disorders and certain site-specific cancers, such as colorectal and breast cancer[1]. Obesity has clinical manifestations such as overweight, excessive body fat and imbalance of metabolism in the body. As the number of obese patients continues to increase, the treatment of obesity becomes a medical problem. In 2010, overweight and obesity caused an estimated 34 million deaths worldwide, which accounting for 3.9% of the total annual mortality rate [2]. In 2017, obesity has the fastest relative growth and became one of the top five risk factors for death and disability [3].

Glucagon-like peptide-1 (GLP-1) is a pro-glucagon secreted by intestinal L cells and pancreatic alpha cells [4]. Turton et al.[5] demonstrated that GLP-1 inhibits food intake through a mechanism mediated by the central nervous system (CNS). There also lots literatures have shown that GLP-1 can promote weight loss and improve lipid metabolism[6].Some studies have shown that systemic administration of GLP-1 or GLP-1 receptor agonists reduces food intake, slows gastric emptying, and reduces body weight [7]. Therefore, GLP-1 has the potential to be developed into an anti-obesity drug, and GLP-1 receptor agonists Liraglutide have been used as anti-obesity drugs for the treatment of obesity.

However, the half-life of GLP-1 is only 2-5 minutes in vivo, which severely limits the biological effects of GLP-1. Therefore, the development of GLP-1 mainly focuses on prolonging the half-life of GLP-1 and promoting its biological effects. A novel GLP-1 derivative was designed in our laboratory in the early stage. LPHSHRAHSLPP was used as the ABD sequence, FNPKTP as the Linker sequence, FNPK as thrombin recognition and restriction site, and TP as dipeptidyl peptidase enzyme hydrolysis site. Then the natural active GLP-1 (7-37) was ligated to obtain a long-acting GLP-1 derivative based on wild-type GLP-1[8]. We evaluated the kinetics and bioactivity of 6-KTP in DIO mice and its long-term effects on food and water intake, body weight, blood lipids, pancreas and fatty liver in the DIO mouse model of obesity.

## 2 Materials and methods

### 2.1 Materials

Long acting GLP-1 derivate 6-KTP was obtained in our molecular biology and biochemistry lab in Jinan University (China).

Normal diet (4% fat, 20% protein) were provided by the Jinan University Experimental Animal Management Center (China) and high fat/sugar diet (40% fat, 20% protein, 40% carbohydrates) were obtained from MaoSibeike biological technology(China).

### 2.2 Animals and treatment

C57BL/6 male mice (8 weeks old) obtained from Jinan Pengyue Experimental Animal Breeding Co. Ltd. were housed in stainless steel hanging cages (four to five per cage) under standard laboratory conditions (12:12 light: dark cycle, temperature 25−27°C, humidity 50%−60%), with free access to food and water. Animal treatment is in accordance with National Institute Health (NIH) guidelines and experimental protocols were approved by the Institutional Animals Care and Use Committee at Jinan University.

All animals were monitored for 2 weeks prior to any experimental procedures. Then the mice were divided in to 2 groups, HFD group (n=30) and the NC group (n=8) were fed with high fat diet and normal diet, respectively. The amount of diet and water was measured every day, and the weight of mice was measured every week to observe the status and weight changes of mice in each group.

Long-term treatment experiment of DIO mice with 6-KTP. Experiments were performed in mature male lean and diet-induced obese (DIO) C57BL/6J mice. The mice were divided in to 4 groups, 8 mice/ group, 6-KTP group were peritoneal injection with 6-KTP every day (1800μg/kg of body weight) on DIO mice, Liraglutide group were peritoneal injection with Liraglutide every day (400μg/kg of body weight) on DIO mice, NC group were peritoneal injection with PBS every day (0.2ml/mice) on lean mice, DIO group were peritoneal injection with PBS every day (0.2ml/mice) on DIO mice for 12 weeks. Food and water intake were measured daily, and the animals were weighed twice per week.

### 2.3 Sample collection and biochemical parameters test

At the end of the administration, the mice in each group were subjected with anesthesia, and blood samples were collected using cryotubes. Serum samples were obtained by centrifugation at 4°C, 1500 g for 20 minutes and then sent to the First Affiliated Hospital of Jinan University for testing. Blood lipids (TG, TC, LDL-C and HDL-C) were analyzed by biochemical auto analyzer. (Wako, Japan).

The adipose tissue were quickly removed, fixed in 4% paraformaldehyde for 24 hours, and then embedded in paraffin. Serial thin sections of 3 μm were sliced and stained with hematoxylin eosin (HE) for histopathology by light microscopy (Olympus BX51 Microscope, Japan).

### 2.4 Data statistical analysis

Student’s t-test or one- or two-way ANOVA were performed to validate statistically significant parameters that are increased or decreased among these two groups. Data processed by Graph Pad Prism software were expressed in mean ± standard error of mean (SEM), and p ≤ 0.05 was considered statistically significant.

## 3 Results

### 3.1 High fat diet increase body weight and food intake

As shown in Fig 1, we confirmed that mice fed with high fat diet could intake more food than that fed with normal diet. The average daily food intake dose of the mice in the HFD group was 39.65 ± 1.998g, and the NC group was 19.93 ± 0.9544g. The high fat diet led to an increase intake in the diet of mice in the HFD group, which was beneficial to lipid accumulation and promoted the construction of the obesity model. (Fig 1)

**Fig 1.**
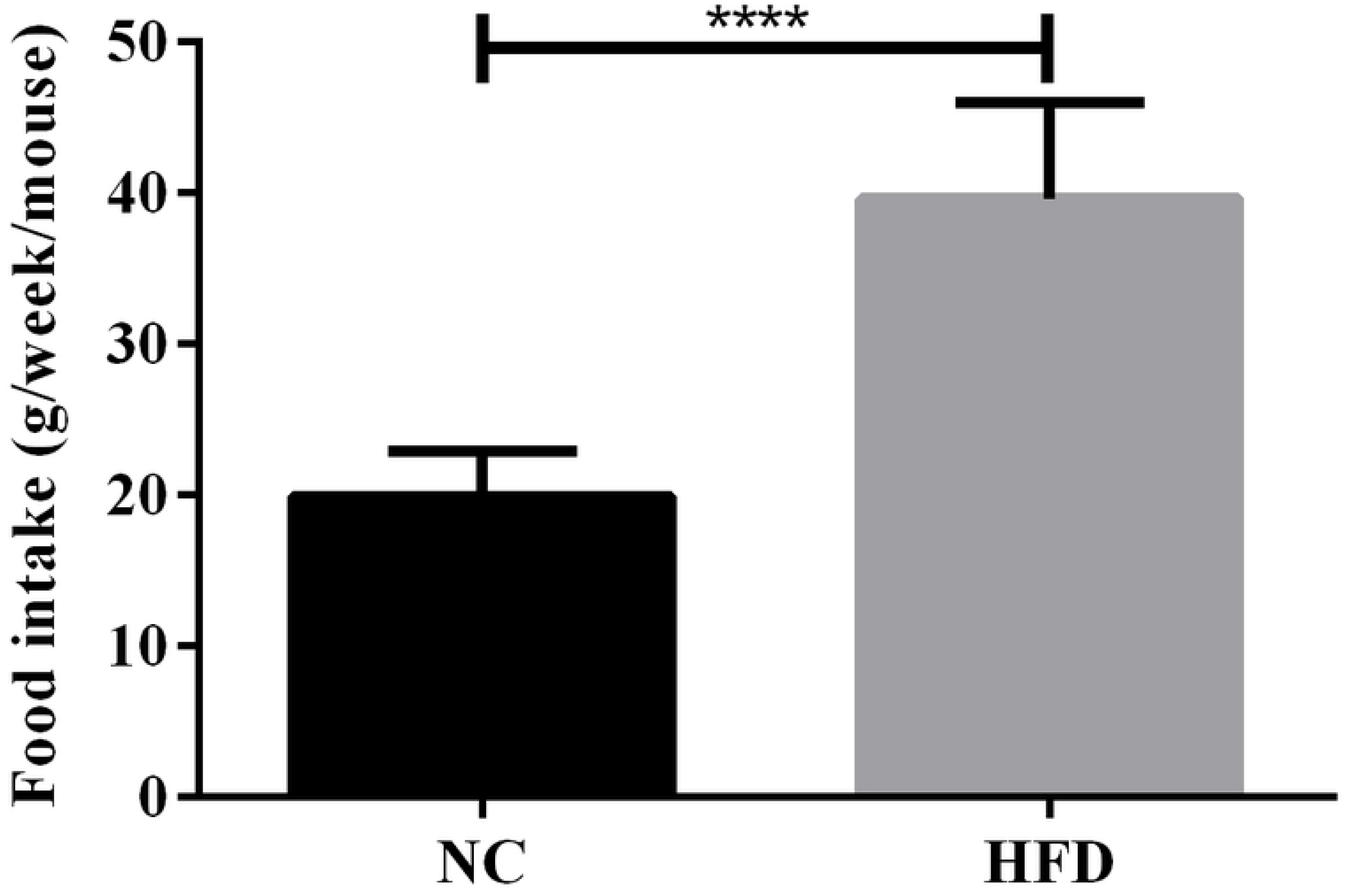
Effect of high-fat diet on diet intake in each group. (*p<0.05, **p<0.01, ***p<0.001, ****p<0.0001)

After 8 weeks, the body weight of the NC group and HFD group were increased. However, as shown in Fig 2, the body weight of the HFD group mice (31.48 ± 0.83g) was significantly higher than NC group mice (25.54 ± 0.61g), approximately 23.25%, reached the obesity model standard. (Fig 2)

**Fig 2.**
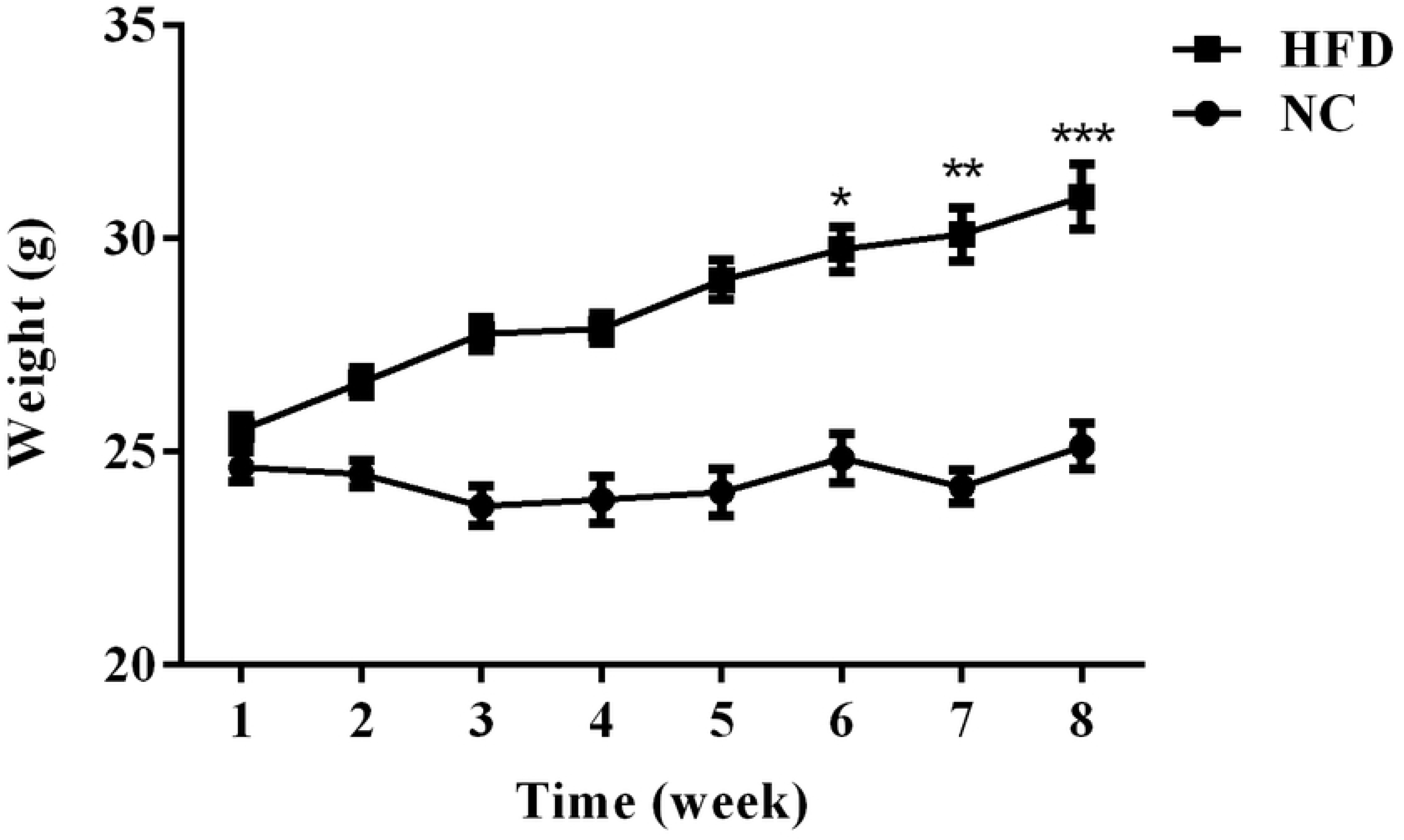
Effect of high-fat diet on body weight in each group. (*p<0.05, **p<0.01, ***p<0.001, ****p<0.0001)

### 3.2 Effect of 6-KTP on body weight and food intake

After a 12-week dosing cycle, statistical analysis of the amount of food intake in each group of mice showed that the long-acting GLP-1 derivative 6-KTP can reduce the food intake of mice. The food intake in the 6-KTP group mice was significantly less than DIO group (p = 0.0197). There was no significant difference between the 6-KTP group and the Liraglutide group (Fig 3).

**Fig 3.**
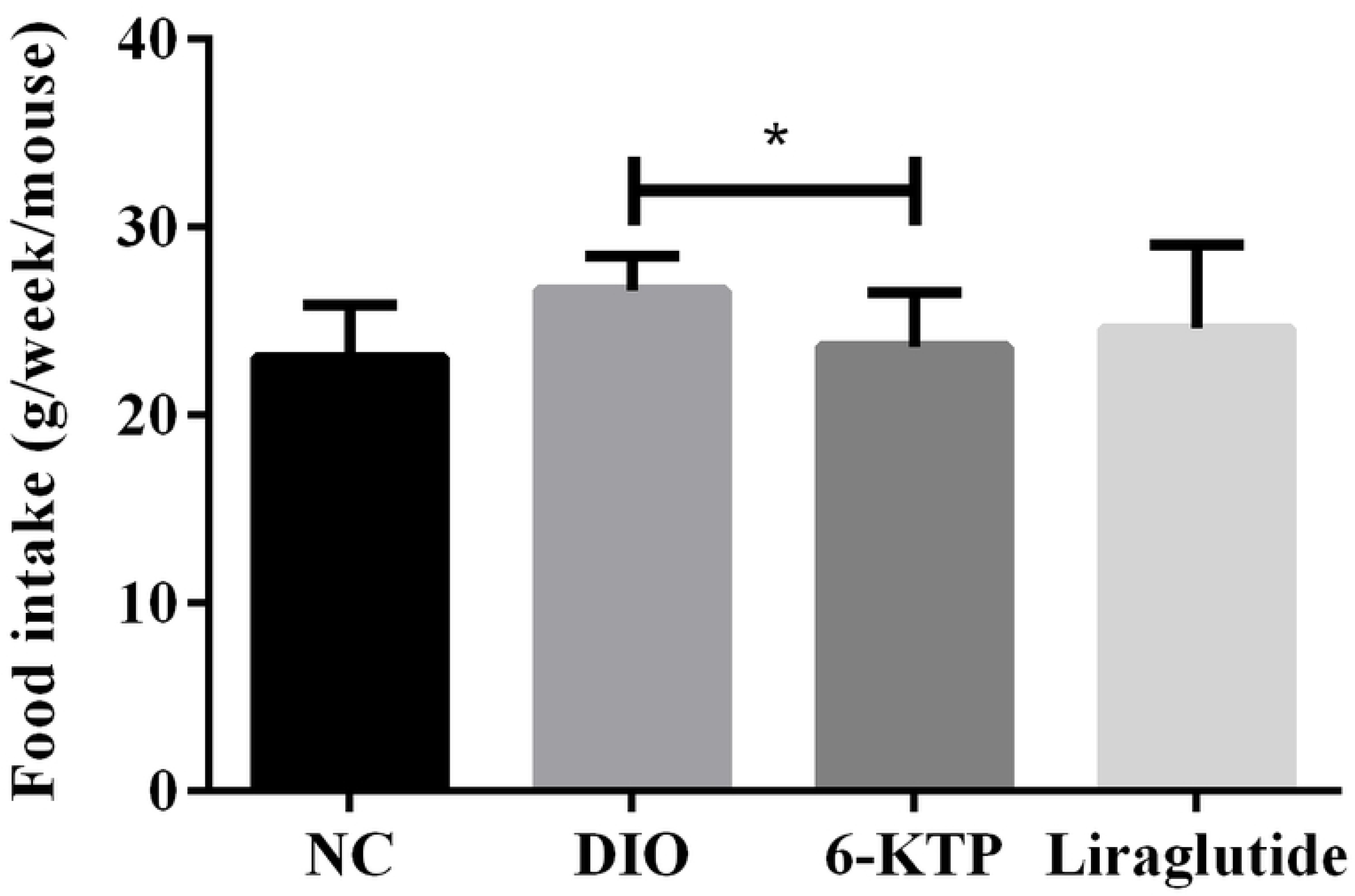
Effect of 6-KTP on food intake in DIO mice.

Repeated once-daily administration of 6-KTP or liraglutide to DIO mice, the body weight was significantly reduced (Fig 4). The body weight in the 6-KTP group mice was significantly less than DIO group (p = 0.0181), whereas, there was no significant difference between the 6-KTP group and the Liraglutide group. The results indicated that 6-KTP had the effect of reducing the body weight of the mice fed with high fat diet.

**Table 1.** Comparison of body weight of mice before and after administration.

| Group | Before treatment (g) | After treatment (g) |
| --- | --- | --- |
| NC group | 26.17±1.57 | 27.91±1.55 |
| DIO group | 31.57±1.88 | 39.30±4.40* |
| 6-KTP group | 32.07±2.37 | 28.71±1.08 |
| Liraglutide group | 31.60±2.26 | 29.64±0.82 |

**Fig 4.**
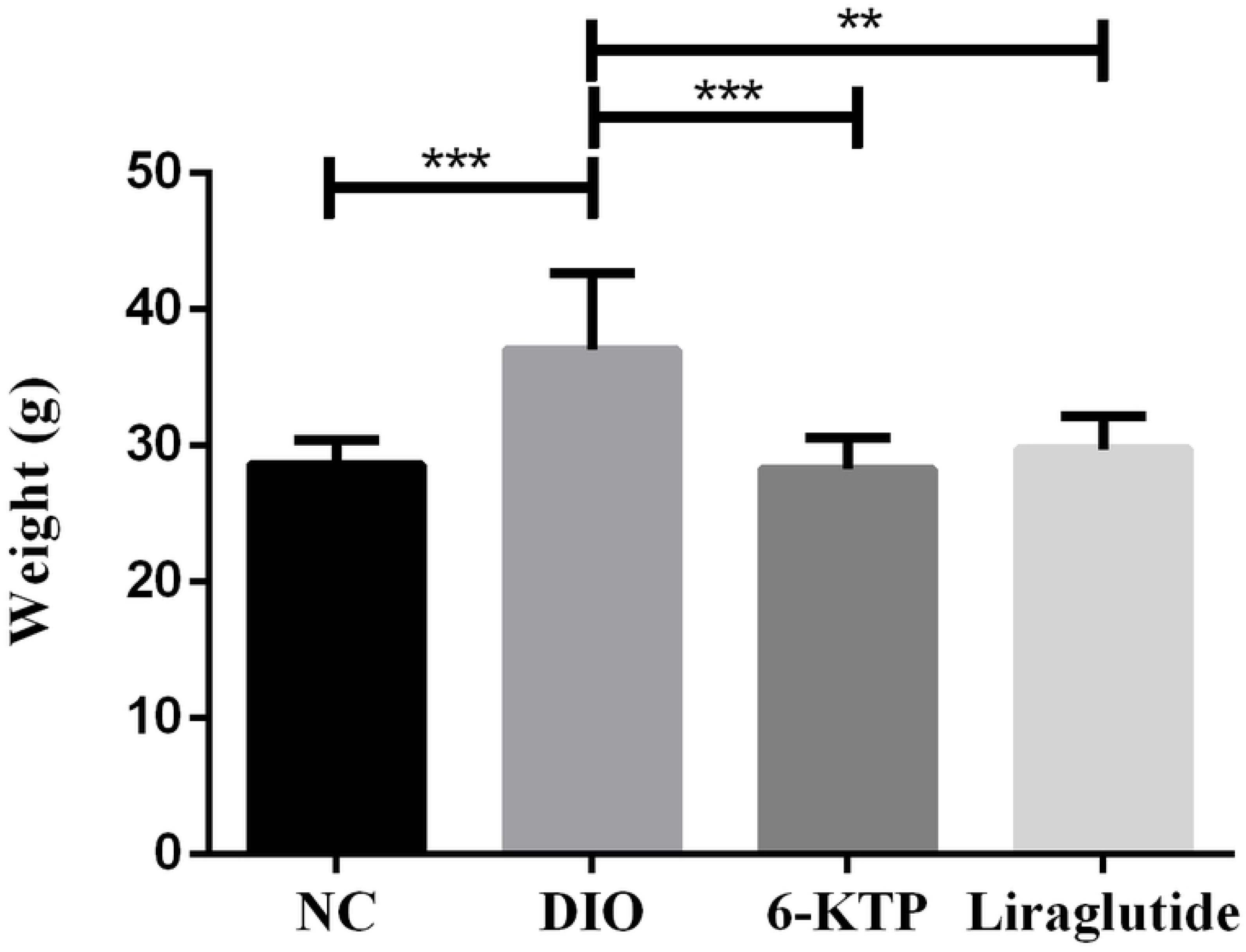
Body weight changes in each group after administration.

### 3.3 Effect of 6-KTP on blood lipid metabolism

To further investigate the potential changes of lipid metabolism, CHOL, TG, HDL-C and LDL-C were analyzed (Fig 5). Compared with the DIO group, the serum levels of Cholesterol (CHOL) were lower than that in the NC group, the 6-KTP group and the Liraglutide group (Fig 5A). There were significant differences between the NC group and the DIO group (p= 0.0008), between the 6-KTP group and the DIO group (p=0.0003), between the Liraglutide group and the DIO group (p=0.0004). There was no significant difference between the other groups. There was no significant difference between the groups of the triglyceride (TG) levels yet (Fig 5B). The levels of serum high density lipoprotein (HDL-C) in the NC group, 6-KTP group and Liraglutide group were lower compared with the DIO group (Fig 5C). There were significant differences between the NC group and the DIO group (p = 0.0006), between 6-KTP group and DIO group (p = 0.0003), between Liraglutide group and DIO group (p = 0.0003). There were no significant differences between other groups. The levels of low-density lipoprotein (LDL-C) in the NC group, the 6-KTP group, and the Liraglutide group were lower than DIO group (Fig 5D). There were significant differences between the NC group and the DIO group (p=0.0160), between the 6-KTP group and the DIO group (p=0.0108), between the Liraglutide group and the DIO group (p=0.0203). There was no significant difference between the 6-KTP group and Liraglutide group. (Fig 5)

**Fig 5.**
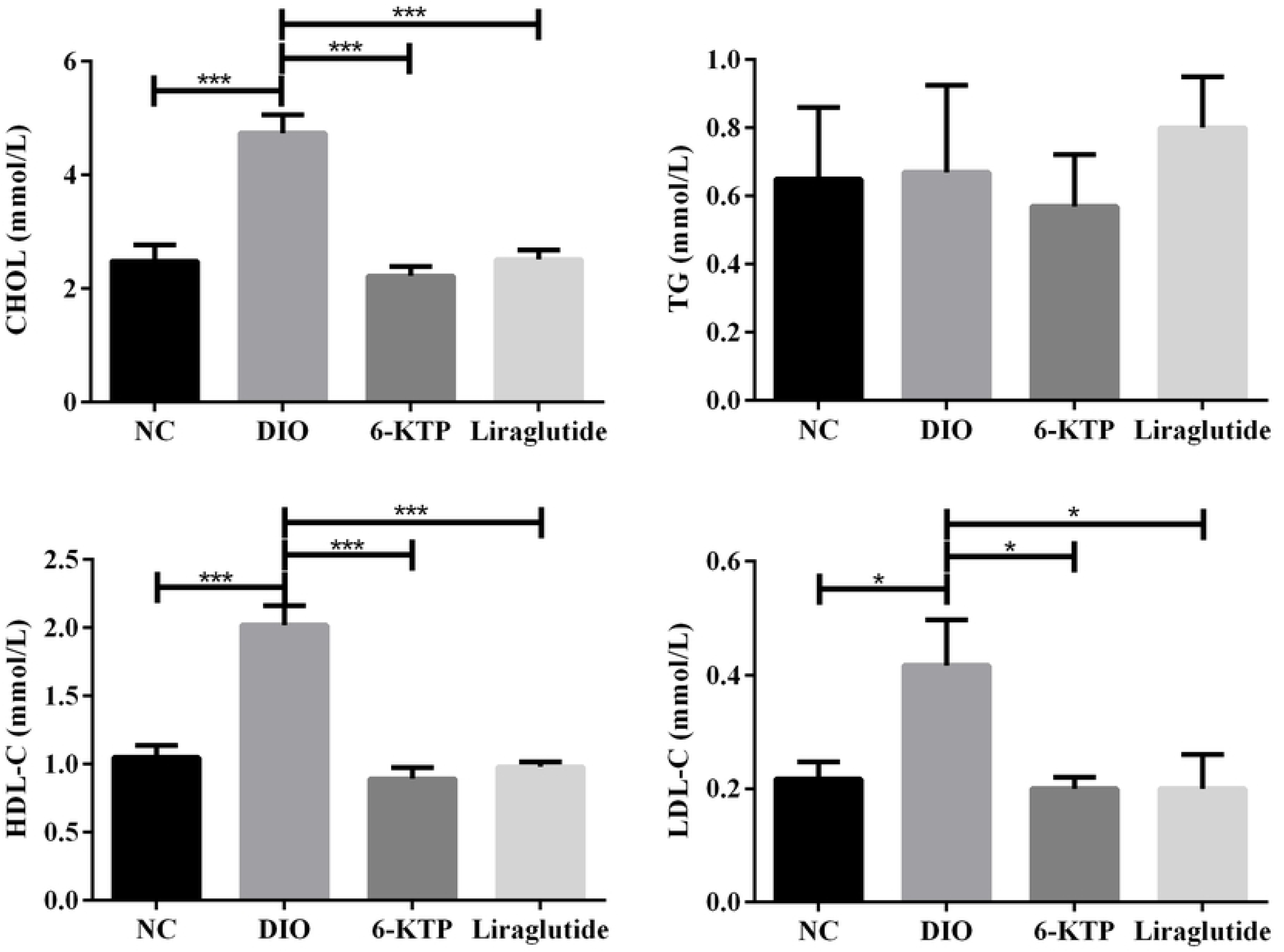
Serum lipid levels in each group of mice. (*p<0.05, **p<0.01, ***p<0.001, ****p<0.0001)

Obvious pathological changes were found in adipose tissue (Fig 6). The DIO group had larger fat cells and more fat accumulation. The fat cells in the 6-KTP group were smaller and had less fat accumulation. There was no significant difference between the 6-KTP group and the Liraglutide group. As shown in Fig 7, the cross-sectional area of the adipocytes in the 6-KTP group was significantly reduced compared with the DIO group, and there was a statistically significant difference (p < 0.0001). There was no significant difference in the cross-sectional area of fat cells in the 6-KTP group compared with the cross-sectional area of the fat cells in the Liraglutide group. (Fig 6,7)

**Fig 6.**
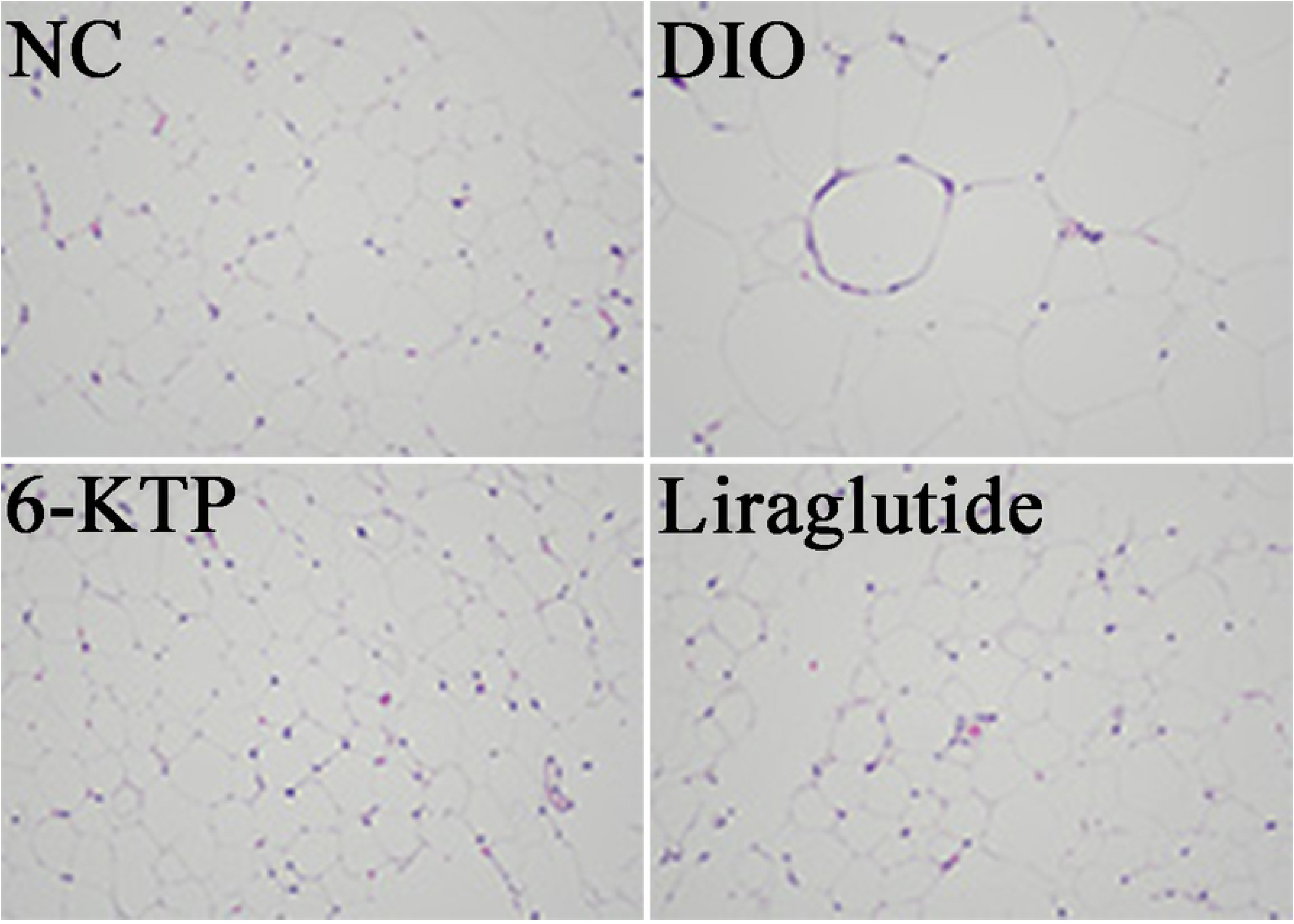
Morphological changes of adipose tissue in each group (H&E, ×200)

**Fig 7.**
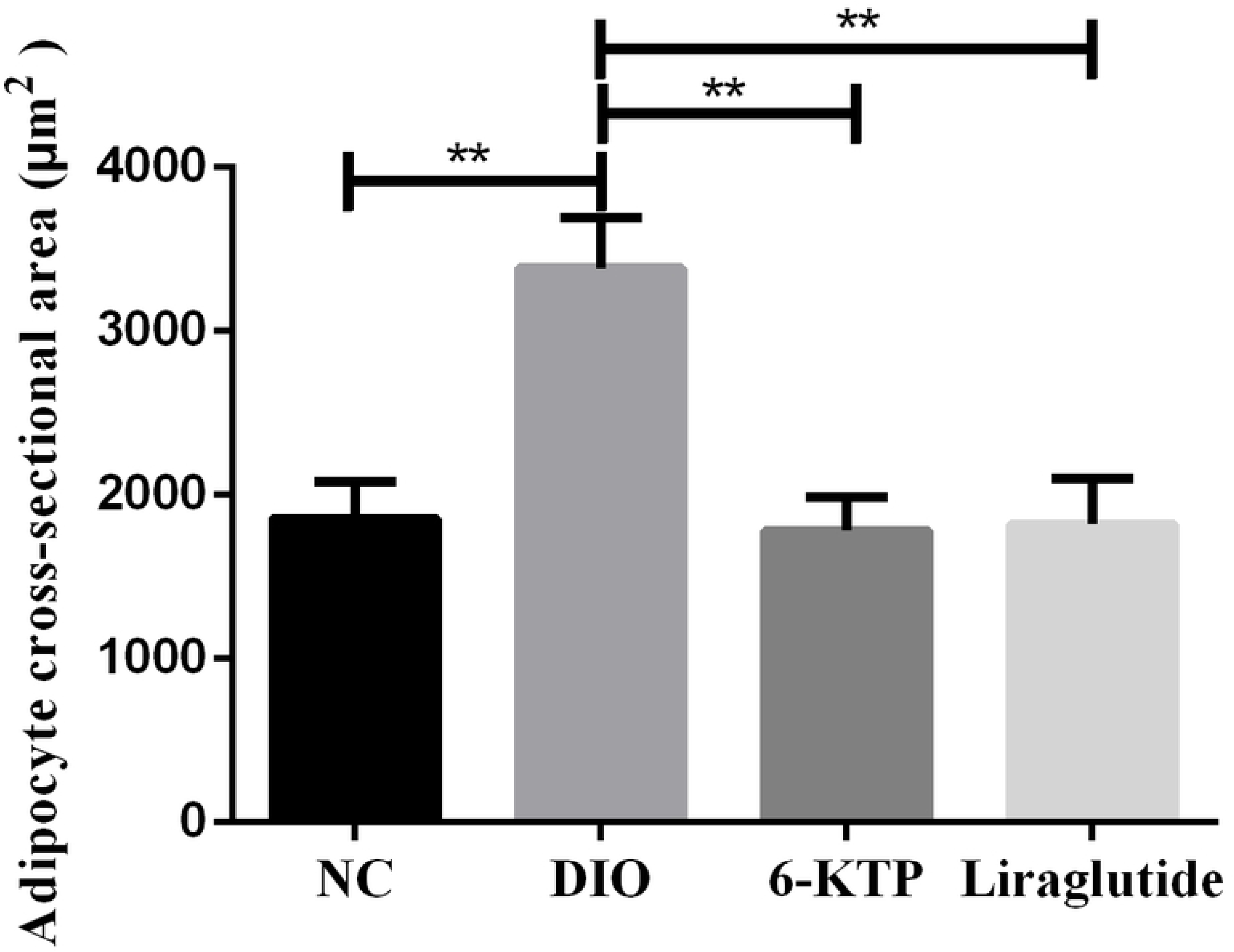
Adipocyte cross-sectional area in each group (μm^2^) (*p<0.05, **p<0.01, ***p<0.001, ****p<0.0001)

## 4 Discussions

Obesity can be divided into three categories according to its causes, including hereditary obesity, secondary obesity and simple obesity. People intake more high fat and high calories food with the improvement of life, which is the main cause of obesity. A large number of studies indicated that people and animals have obesity susceptibility under high-fat diet conditions [9]. A study found that high fat diet induced obesity caused resistance to insulin signaling in the major PI3K and ERK signaling pathways, and the results suggest that the process is mediated by inflammation within BAT itself.

It has been shown[10] that GLP-1 has an effect on increasing satiety and suppressing appetite. GLP-1 exerts this effect through various pathways. In rodents, intraventricular injection of GLP-1 can reduce food intake. In human patients, injection of GLP-1 can also increase satiety and reduce energy intake. Many ring-shaped organs, including the infraorbital organs, organ blood vessels at the end of the lamina, median bulges, posterior pituitary and posterior regions, contain high-density GLP-1 binding sites[11]. These sites may function as a window of the brain, and the peripherally released GLP-1 affects the CNS through this pathway. Our results also show that the long-acting GLP-1 derivative 6-KTP has an effect of suppressing appetite and reducing the amount of food.

This is consistent with the literature[12], animal model tests show that GLP-1 drug Liraglutide mainly reduces body weight by reducing energy intake mechanism, may also change food preferences and maintain energy consumption to reduce body weight. The results showed that the long-acting GLP-1 drug 6-KTP can inhibit the weight gain of the high fat diet-induced obese mouse model and promote its weight loss. Compared with the positive drug Liraglutide, there was no significant difference in body weight between the two groups. Compared with the DIO group, the mice after 6-KTP treatment showed a significant decrease in body weight, and there was a statistically significant difference.

It has been reported in the literature [13], that Liraglutide has an effect on improving the levels of CHOL and LDL-C in patients. The results showed that after treatment with long-acting GLP-1 drug 6-KTP, the mice were treated with TG, CHOL and HDL-C and LDL-C levels have improved. The levels of CHOL, HDL-C and LDL-C were statistically different compared with the DIO group, and the TG levels were decreased but not statistically different. The four levels of blood lipids in the long-acting GLP-1 drug group 6-KTP group were not significantly different from those in the Liraglutide group, indicating that 6-KTP has the same effect as the positive drug. By comparing the four levels of blood lipids in the 6-KTP group and the DIO group, the results showed that 6-KTP can effectively improve the lipid metabolism disorder caused by the high-fat diet, promote the recovery of lipid metabolism, inhibit the abnormal accumulation of lipids, and thus achieve weight inhibition.

In obesity, white adipose tissue may become dysfunctional and thus unable to expand properly to store excess energy, while at the systemic level, these dysfunctions lead to other tissues that regulate metabolic homeostasis (e.g. liver and endocrine pancreas) Ectopic fat deposition in the middle. This will lead to other diseases such as: insulin resistance progression and increased risk of T2DM[14]. The results showed that the 6-KTP group adipose cells became smaller and adipose accumulation was reduced. There was no significant difference between the Liraglutide and the 6-KTP. 6-KTP can inhibit fat accumulation and promote adipose degradation, thereby promoting weight loss. In one study, liraglutide, a GLP-1R agonist administered, stimulated brown adipose tissue (BAT) heat production and adipocyte browning, and the mechanism controlling these effects was located in the hypothalamic ventromedial nucleus (VMH)[15]. The mechanism controlling these actions is located in the hypothalamic ventromedial nucleus (VMH), and the activation of AMPK in this area is sufficient to blunt both central liraglutide-induced thermogenesis and adipocyte browning.

## 5 Conclusions

Our results indicated that the long-acting GLP-1 derivative 6-KTP can inhibit the increase of body weight in mice by reducing the amount of food. Furthermore, under GLP-1 drug treatment, there was no difference in the food intake and body weight gain in Liraglutide group. 6-KTP has the effect of promoting lipid metabolism and inhibiting fat accumulation, and there was no significant difference between the results and the liraglutide group. The finding suggested the GLP-1 derivate 6-KTP may therefore affect the development of obesity and may be a promising new treatment in patients with this condition.

